# Mechanisms of phosphatidylserine influence on viral production: a computational model of Ebola virus matrix protein assembly

**DOI:** 10.1101/2021.07.22.453424

**Authors:** Xiao Liu, Ethan J. Pappas, Monica L. Husby, Balindile B. Motsa, Robert V. Stahelin, Elsje Pienaar

**Author notes:** Co-first authors (Xiao Liu, Ethan J. Pappas). Indiana University School of Medicine (Ethan J. Pappas).

## Abstract

Ebola virus (EBOV) infections continue to pose a global public health threat, with high mortality rates and sporadic outbreaks in Central and Western Africa. A quantitative understanding of the key processes driving EBOV assembly and budding could provide valuable insights to inform drug development. Here we used a computational model to evaluate EBOV matrix assembly. Our model focused on the assembly kinetics of VP40, the matrix protein in EBOV, and its interaction with phosphatidylserine (PS) in the host cell membrane. Human cells transfected with VP40-expressing plasmids are capable of producing virus-like particles (VLPs) that closely resemble EBOV virions. We used data from this in vitro VP40 system to calibrate our computational model. PS levels in the host cell membrane had been shown to affect VP40 dynamics as well as VLP production through recruiting VP40 dimers to plasma membrane inner leaflet. Our computational results indicated that PS may have direct influence on VP40 filament growth and affect multiple steps in the assembly and budding of VP40 VLPs. We also proposed that the assembly of VP40 filaments may follow the nucleation-elongation theory where initialization and oligomerization of VP40 are two separate and distinct steps in the assembly process. This work illustrated how computational and experimental approaches can be combined to allow for additional analysis and hypothesis generation. Our findings advanced understanding of the molecular process of EBOV assembly and budding processes and may help the development of new EBOV treatments targeting VP40 matrix assembly.

## Introduction

EBOV was first identified in 1976 with two different outbreaks in Africa, where 284 and 318 people were infected with 53% and 88% mortality, respectively (1, 2). Since then, almost 30 known EBOV outbreaks have occurred, together causing more than 30,000 cases and 13,000 deaths (3). Recent outbreaks in Uganda, Guinea and the Democratic Republic of Congo illustrate the continued threat from this deadly infection.

Some therapies have shown promise as treatment for Ebola virus disease (EVD) in pre-clinical animal models (4–9). Experimental treatments were also tested in humans during the multi-country outbreak in 2014-2016 including monoclonal antibody cocktails (10, 11), anti-viral drugs (12–15) and other therapies (16–22). Antibody cocktails and anti-viral drugs have also been administered in the most recent EBOV epidemic in the Democratic Republic of Congo and, together with supportive care, reduced fatality rates (23, 24). However, most of these investigational treatments were given as Monitored Emergency Use of Unregistered and Investigational Interventions (MEURI) (23). Only two monoclonal antibody therapies were recently approved for the treatment of EVD, but mortality remains high even with these treatments (more than 30%), and sideeffects can be severe (25–27). Developing effective and safe EBOV therapies is challenging, in part, due to our limited understanding of key mechanisms of protein-protein and lipid-protein interactions in the EBOV life cycle.

EBOV pathogenesis studies are challenging because EBOV research is limited to facilities with BSL-4 infrastructure. In order to study parts of the EBOV life cycle in lower safety level laboratories, different viral genes have been inserted into plasmids and expressed separately or together in transfected cells. Matrix protein VP40 (VP40) is the main component of the EBOV matrix and is critical for EBOV assembly and budding. VP40, when expressed independently of the other six EBOV proteins, has been shown to form VLPs with similar size, shape, cell attachment and entry properties as EBOV virions (28–31). These non-infectious VP40 VLPs therefore represent a useful tool for studying EBOV assembly and budding processes *in vitro* in BSL-2 conditions.

This VP40 system has generated key insights into VP40 assembly. VP40 was shown to form homodimers in the cytoplasm through N-terminal domain interactions, abrogation of which halts plasma membrane localization and budding of VP40 (32). VP40 dimers bind to the cell membrane through interactions between VP40 C-terminal domains and PS (33). VP40 membrane dimers further assemble into hexamers and larger oligomers in the growing virus filaments (32, 34). There is evidence that the assembly and budding of VP40 VLPs is dependent on the level of PS in the host cell membrane (33, 35), suggesting that PS levels or VP40-PS interactions could be a drug target to disrupt EBOV reproduction. Membrane PS levels have been shown to affect VP40 membrane binding (35) and PS levels also impact the relative number of VP40 oligomers and VLPs produced over 48 hours (35, 36). However, it remains unclear if the impact of PS on VP40 membrane association is able to account for the observed impacts on VP40 oligomer levels and VLP production (Fig. 1). Furthermore, the dynamics of VP40 filament growth has not been reported, making it difficult to predict the longterm impacts of disrupting VLP assembly by targeting PS. These questions are critical for the development of EBOV treatment, but difficult to answer using experimental approaches alone. While VP40 structural information and lipid binding data at the membrane are available, it is not always possible to experimentally uncouple individual steps in the VLP assembly process (e.g., VP40 membrane binding and oligomerization), or to obtain high temporal- and spatial-resolution data to quantify VP40 dynamics.

**Figure 1.**
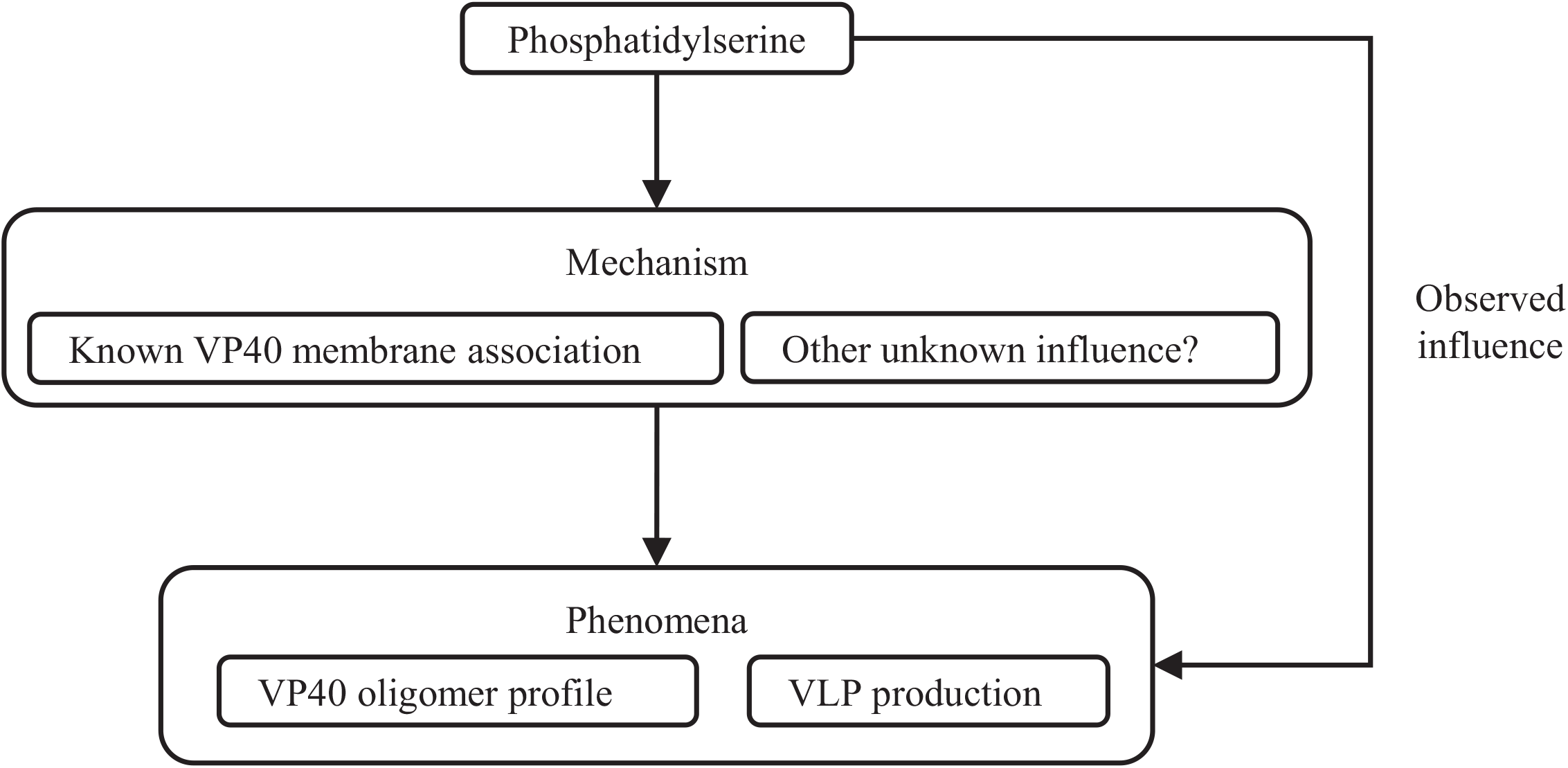
The influence of PS on the EBOV budding process. PS is known to influence VP40 dimer membrane association, oligomer profile and VLP production. However, whether the observed phenomena can be explained by the VP40 dimer association alone remains unclear.

Computational models offer complementary strengths to experimental approaches. Computational models have been applied to evaluate viral infection dynamics in populations (37, 38), physiological disease progression and treatment efficacy (39, 40), as well as viral replication, cellular immune responses (41, 42). For EBOV, computational models have been applied on both population (43–45) and physiological (40, 46–49) levels. In contrast, subcellular models for EBOV assembly are not currently available. However, emerging experimental data and a more detailed understanding of the EBOV replication cycle (35, 36) create an opportunity for a quantitative, computational characterization of EBOV assembly.

In this study, our goal was to quantify the impact of PS in individual steps in the VP40 assembly process and investigate how these steps contribute (or not) to altered VLP production. We developed a subcellular-level ordinary differential equation-based (ODE-based) model of EBOV VP40 assembly and budding. We built and calibrated the model using experimental data from VP40 studies (35, 50), and the model reproduced both qualitative and quantitative experimental observations. We applied our model to evaluate the dynamics of VP40 oligomer accumulation and filament growth, as well as the impact of PS on these processes. Our work showed how computational modeling can facilitate data integration, analysis and interpretation Furthermore, our findings advanced understanding of EBOV assembly and could inform development of PS-targeted treatments of EVD in the future.

## Results

### A novel ODE-based model of EBOV VP40 assembly and budding process

Our computational model spanned VP40 monomer production to VLP budding (Fig. 2). VP40 monomers were produced, dimerized and bound to the host cell membrane. Membrane bound VP40 dimers further combined to form hexamers. Hexamers served as the building blocks for longer VP40 oligomers which form filaments. Fully developed filaments bud to form VLPs. Throughout this paper we explored the possible influence of PS levels on mechanisms throughout the assembly and budding process including membrane binding of VP40 dimers, hexamer formation, oligomerization, filament stabilization and VLP budding.

**Figure 2.**
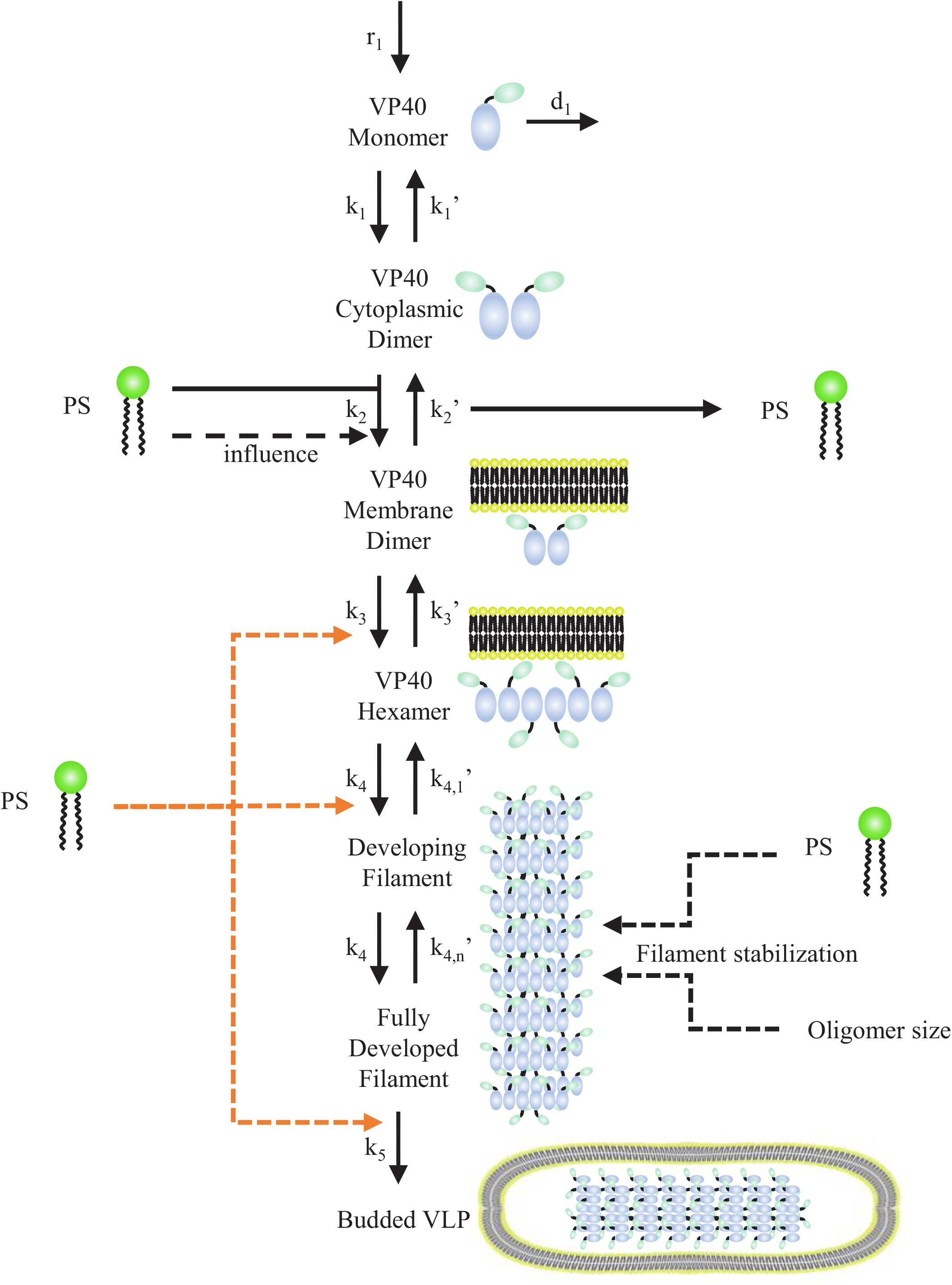
VP40 ODE-Model Structure. The model starts with production of VP40 monomer and ends with production of budded VLP. The impacts of PS on membrane association (k_2_), hexamer formation (k_3_), oligomerization (k_4_, k_4,n_’), and VLP budding (k_5_) are shown in dashed lines. The orange dashed lines are three influences of PS evaluated separately in our extended models (Ex1, Ex2 and Ex3). Black dashed lines are included in the baseline model and all extended models.

The experimental data that we aimed to reproduce are described in detail in Methods, but the key observations that were used to evaluate different PS mechanisms are summarized here:

- Lower PS levels were associated with lower oligomer ratio (defined as the ratio of VP40 hexamers or larger oligomers over dimers and monomers (35))
- Transfected cells produced approximately 1×105 VLPs 24h after transfection.
- The relative frequency of oligomers decreased from hexamers to 42-mers, and this decrease was more pronounced at lower PS levels.
- VLP production was reduced at lower PS levels, and this reduction was sustained over 48h after transfection.

Our initial analysis aimed to determine if the known impact of PS on VP40 membrane association was enough to produce the observed changes in VP40 oligomer levels and VLP production (Fig. 1). Our preliminary model (Table 1) therefore only included the known influence of PS level on the dissociation constant (K_D2_) of VP40 dimer binding to cell membrane (33, 35). Our results indicated that this preliminary model was unable to reproduce the experimentally measured impact of PS on VLP production and oligomer frequencies simultaneously. The experimentally observed decrease in oligomer frequencies from hexamer to 42-mer can only be captured in the model when most VP40 have not bound to membrane, leading to no detectable VLP production (Fig. S1A). Conversely, when VLP production can be observed, the frequencies from hexamer to 42-mer become very similar (Fig. S1B).

**Table 1.** Model construction.

| Included Mechanism | Model |  |  |  |  |
| --- | --- | --- | --- | --- | --- |
|  | Preliminary | Baseline | Ex1 | Ex2 | Ex3 |
| PS influence on VP40 membrane association | √ | √ | √ | √ | √ |
| Filament stabilization |  | √ | √ | √ | √ |
| PS influence on filament stabilization |  | √ | √ | √ | √ |
| PS influence on hexamer formation |  |  | √ |  |  |
| PS influence on filament growth |  |  |  | √ |  |
| PS influence on budding rate |  |  |  |  | √ |

These findings suggested that PS impact on VP40 dimer membrane association alone is not able to explain observed PS impacts on VLP production and oligomer frequencies. To identify possible unknown mechanisms in the assembly and budding process of VP40, we built four sub-models (Baseline and three extended models: Ex1, Ex2, and Ex3) (Table 1). In subsequent sections we will describe our progressive addition of unknown mechanisms, and integration of experimental datasets.

### Filament stabilization is necessary to reproduce experimentally measured relative oligomer frequencies

To address the inability of the preliminary model to match experimental results for both VLP production and oligomer frequencies, we introduced a filament stabilization mechanism.

We hypothesized that the physical structure of a growing filament and an oligomer consisting of only a few hexamers are very different. We therefore proposed that the structure for filament became more stable as the oligomers become larger. Mathematically we represented this “stabilization” mechanism for the growing filament by decreasing the reverse rate constant of oligomerization (k_4,n_’, Fig. 2) as the oligomer grows. Addition of this stabilization step enables the model to reproduce both the decreasing oligomer frequencies and VLP production. However, without direct influence of PS on the stabilization step, the model was unable to reproduce the experimental differences in oligomer frequencies among PS groups. These results suggested that PS level can have a direct influence on filament stabilization.

We therefore constructed our baseline model (Table 1) that included filament stabilization as well as direct PS influence on this stabilization. The baseline model successfully reproduced the difference in oligomer ratio between the low and high PS groups (Fig. 3A). Furthermore, the baseline model reproduced relative oligomer frequency and VLP production simultaneously (Fig. 3B, D). Difference in relative oligomer frequency among PS groups could also be observed in our simulation (Fig. 3D). The model slightly overestimated VP40 budding ratio compared to experimental data (Fig. 3C).

**Figure 3.**
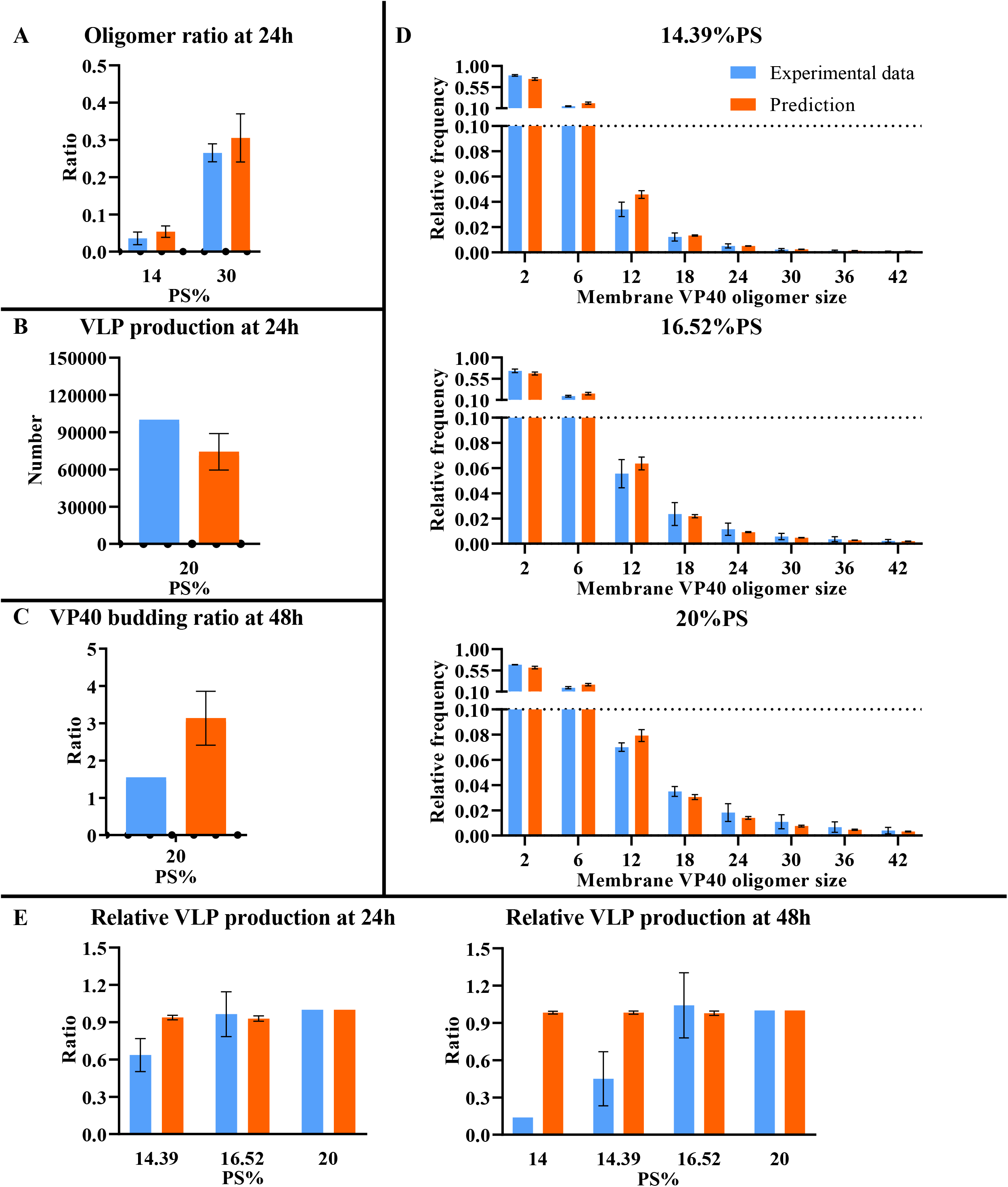
Comparison between computational results from the baseline model and experimental data. Model fit is evaluated based on differences between simulated and experimentally measured: (A) Oligomer ratio at 24 h. (B) VLP production at 24 h. (C) VP40 budding ratio at 48 h. (D) Oligomer frequency at 24 h for 3 different PS levels (14.39%, 16.52%, and 20%). (E) Relative VLP production at 24 and 48 h. Error bar indicates the standard error of the mean (SEM). Simulation data represents top 5 fits. Sample sizes of each experimental data are shown in Table S13.

Considering VLP assembly dynamics, our baseline model predicted that the concentrations of monomer and dimer were increasing toward a steady state over time, while concentrations of hexamer and higher oligomers showed fluctuations. Our model further indicated that dimer was the predominant form of VP40 in cytoplasm (Fig. 4A, S2), which is aligned with experimental observations (51). Although the concentration of cytoplasmic VP40 (monomer and dimer) was much higher than that of VP40 oligomers on the cell membrane, the overall amount of VP40 bound to the membrane was higher than the amount of VP40 in the cytoplasm due to the size and number of the oligomers (Fig. 4B). This is also aligned with experimental observations (35). These observations provided qualitative validation of our model predictions.

**Figure 4.**
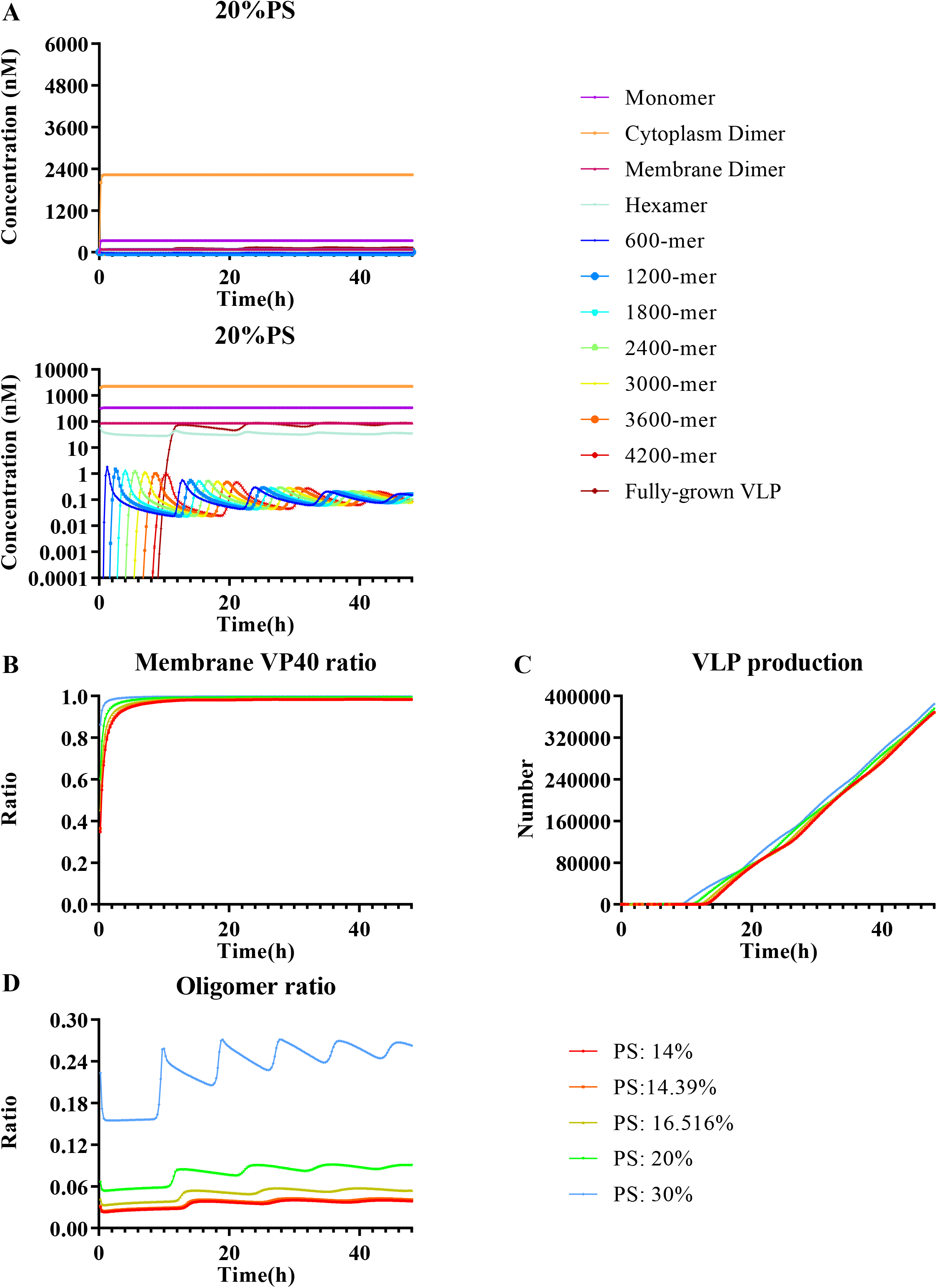
Model predicted system dynamics of best fit in the baseline model. Using the calibrated baseline model, we predict: (A) Time course of VP40 monomer and oligomers. The upper sub-panel is on linear-scale and the lower sub-panel is on log-scale. (B) Time course of membrane/cytoplasmic VP40 ratio. (C) Time course of VLP production. (D) Time course of Oligomer ratio.

We next shifted focus from the subcellular VP40 dynamics to overall VLP production dynamics. The baseline model was unable to reproduce differences in VLP budding among PS groups (Fig. 4C). In contrast, oligomer ratio was positively corelated to PS level (Fig. 4D). The reason is that the number of membrane VP40 oligomers did not dramatically change among different PS groups, while the number of cytoplasmic VP40 monomers and dimers varied significantly (Fig. S2). As a result, oligomer ratio varied among PS groups, while VLP production remained identical. As a result, our baseline model failed to reproduce the experimentally measured relative VLP production among PS groups at 24 and 48 hours (Fig. 3E). Model predictions showed either identical VLP production among PS groups or differences due to fluctuations in time (Fig. S3). However, the impact of fluctuation decreased with time. No calibrated parameter sets for the baseline model could maintain the difference in VLP production among PS groups for 48 hours. Taken together, these results indicated that filament stabilization contributes to progressively decreasing oligomer frequencies in VP40 VLPs, and that this mechanism is required for successful VLP budding. Further, our results suggested that there are other PS influences on VP40 assembly, not included in the baseline model, which drive the observed difference in VLP production among PS groups.

### Direct PS effect on VLP budding rate is required to reproduce experimentally measured longer term differences in VLP production rates

In the work of Adu-Gyamfi et al. (35), PS levels were shown to affect assembly and budding of VLPs. Our baseline model further suggests that the mechanisms by which PS impacts assembly and budding may not be limited to VP40 membrane binding, but also extend to steps downstream of membrane binding. To test this possibility, we built three additional extended models that include influence of PS on VP40 hexamer formation (k_3_, Ex1), filament growth (k_4_, Ex2) and VLP budding (k_5_, Ex3). We calibrated each of these models independently to assess their ability to reproduce the relative VLP production at 24 and 48 hours.

When PS affected hexamer formation (k_3_, Ex1) or filament growth (k_4_, Ex2), the difference among PS levels remained small and mainly depended on fluctuation and the time point where VLPs started budding (Fig. S4, S5). No longer term (48h) effect on relative VLP production was observed (Fig. S6, S7). In contrast, when VLP budding (k_5_, Ex3) was being affected, consistent differences could be observed over long periods for each VLP budding curve (Fig. 5). This Ex3 model was the only one that could match experimentally measured relative VLP production (Fig. 6E), while not losing accuracy in other predictions (Fig. 6A-D).

**Figure 5.**
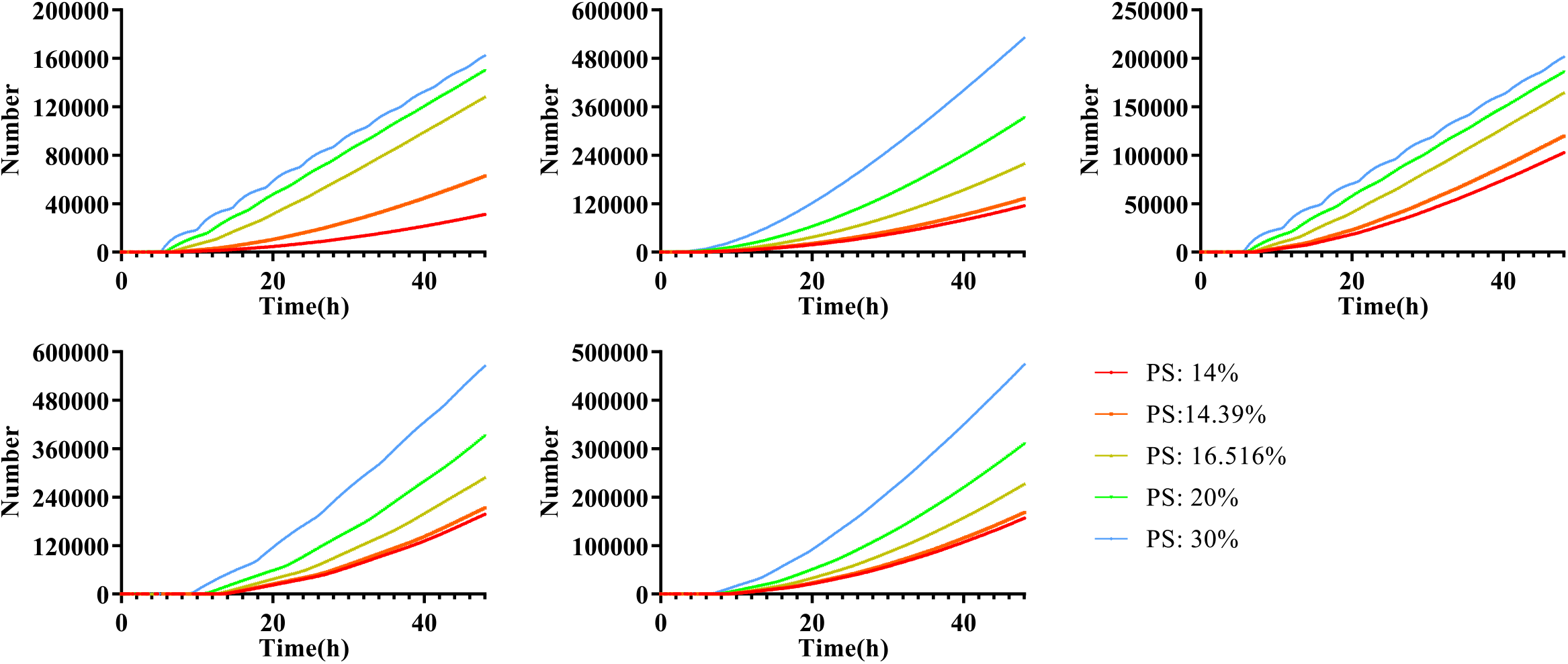
Model predicted VLP production dynamics of the Ex3 model. Each panel represents one of the top 5 model fits, and colors represent different PS levels. VLP production for each PS concentration using the Ex3 model are separated and different.

**Figure 6.**
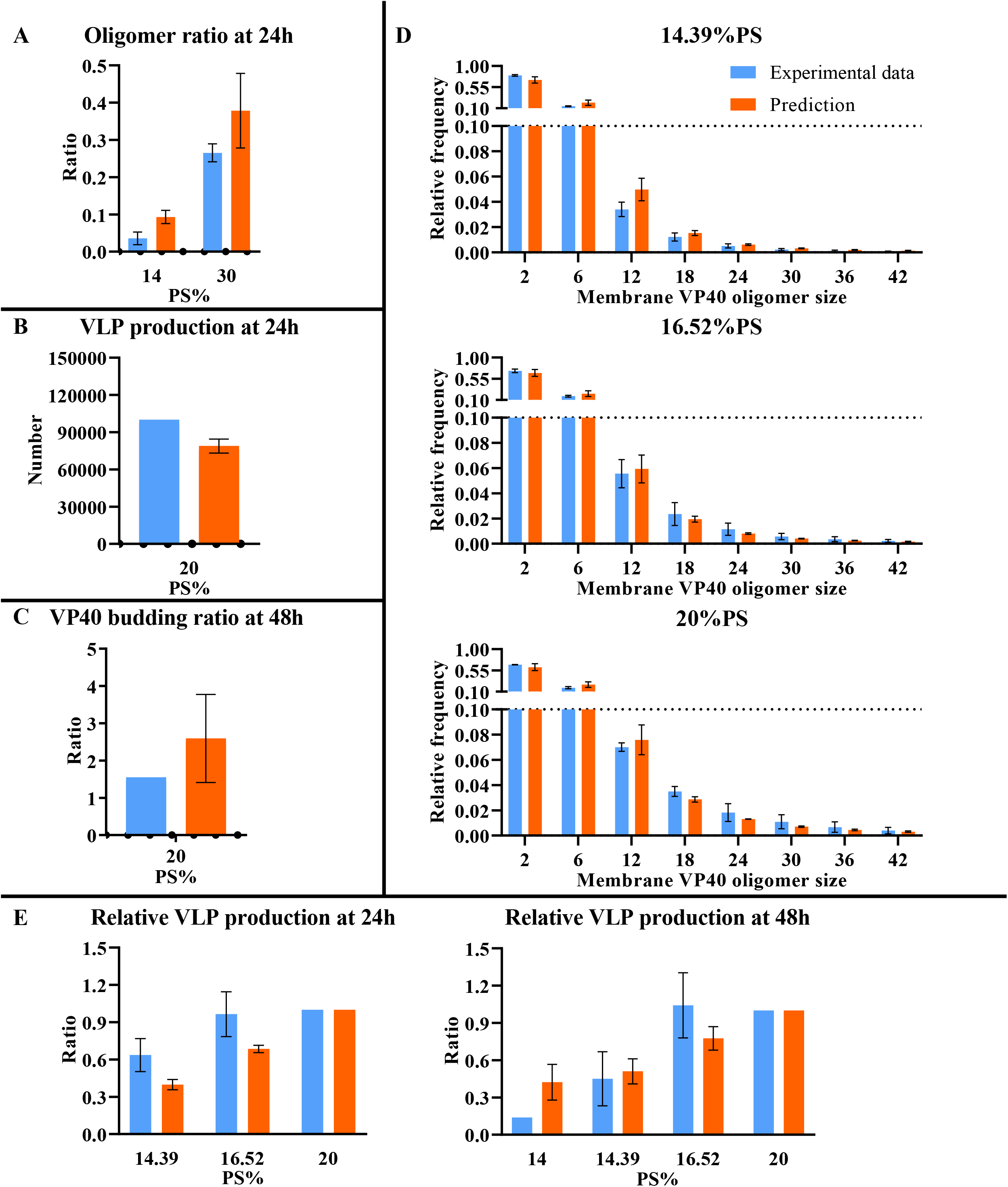
Comparison between computational results from the Ex3 model and experimental data. Model fit is evaluated based on differences between simulated and experimentally measured: (A) Oligomer ratio at 24 h. (B) VLP production at 24 h. (C) VP40 budding ratio at 48 h. (D) Oligomer frequency at 24 h for 3 different PS levels (14.39%, 16.52%, and 20%). (E) Relative VLP production at 24 and 48 h. Error bar indicates the SEM. Simulation data represents top 5 fits. Sample sizes of each experimental data are shown in Table S13.

To quantitatively compare how well each extended model matched all of the experimental data, we chose the top 5 parameter sets for each model and compared the average cost values. The average cost for Ex3 model was the lowest (21.97, Fig. 7, Table S1, S3) while the averages for the other 3 models were similar.

**Figure 7.**
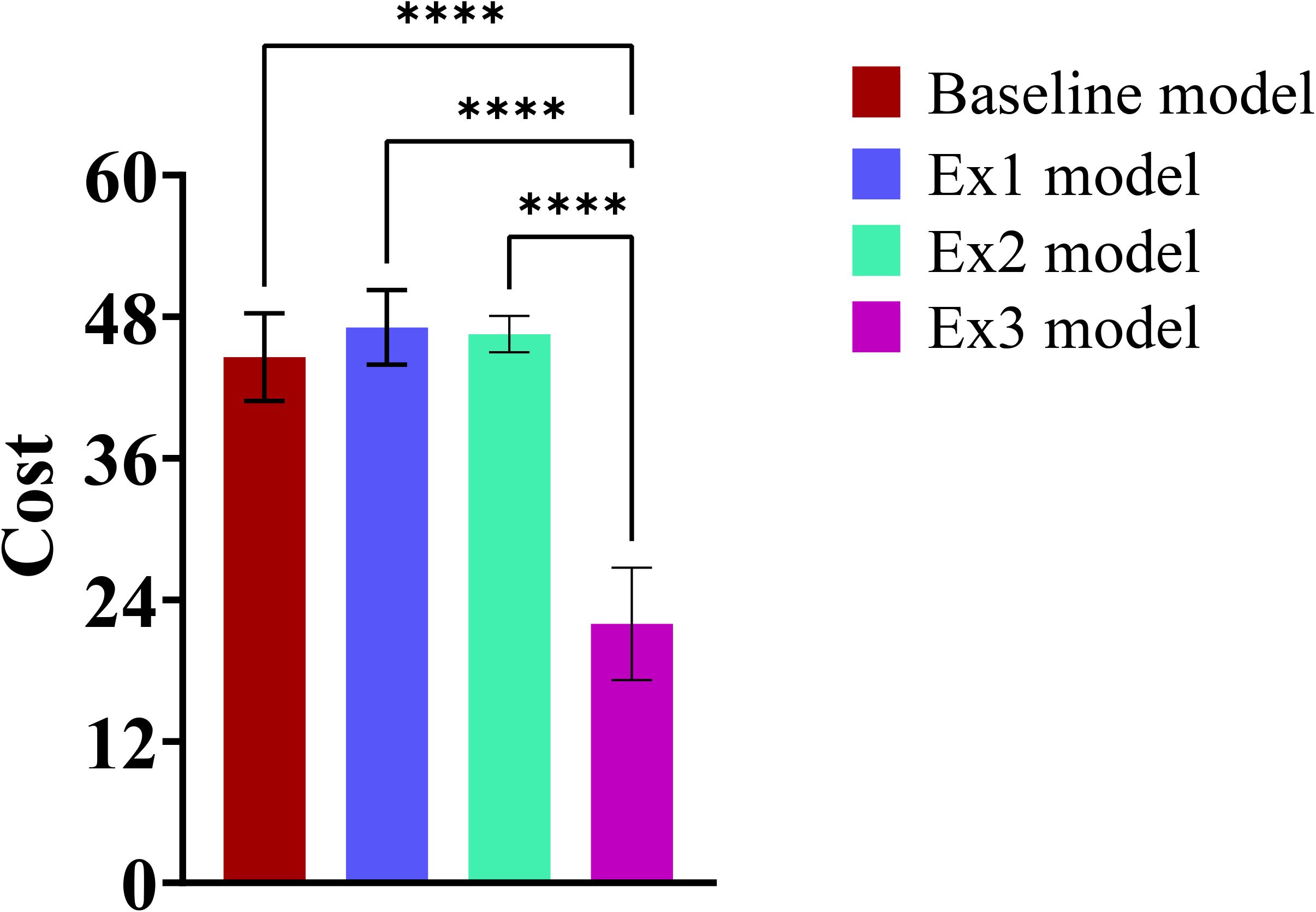
Cost of the top 5 best fits for all models (Baseline, Ex1, Ex2, Ex3). Costs are compared among models using ANOVA and there are statistically significant differences (p-value <0.001) in cost among those models (Table S2). Least Significant Difference (LSD) was conducted to determine statistically significant pairwise differences, and statistical differences are identified between Ex3 and other models (Table S4).

All of our models assumed that VP40 hexamers were the buildings unit of VP40 assembly (28, 29, 35, 36, 50, 52–54). Recent work indicated that the building block for VP40 VLPs can be dimers instead of hexamers(34). To assess the impact of this hypothesis, we modified our Ex3 model to represent a “dimer assembly” system (Fig. S8). The predictions were similar to our hexamer assembly Ex3 model (Fig. S9), indicating that our conclusions were independent of the specific building block of VP40 VLPs.

To validate our predictions before 24 h, we compare our simulations to experimentally measured VP40 membrane localization at both 8h and 24h. Though there are some differences between the absolute values of the simulations and experimental data (Fig. S10A), our simulations correctly predict that the membrane localization is nearly identical between 8 h and 24 h (Fig. S10B). Original data for membrane localization experiments are given in Table S18 and S19.

Thus, our analysis of models Ex1-Ex3 suggested that PS has direct influence on mature filaments budding from the plasma membrane, and that this interaction is important for the observed impacts of PS levels on VLP production.

### Sensitivity Analysis identified key mechanisms that can inform treatment development

To quantify the contribution of individual steps in VP40 assembly to VLP production, the difference among sub models and the influence of PS on the system, we performed global sensitivity analysis using Partial Rank Correlation Coefficients (PRCC). The relationship between 7 parameters (k_1_, k_2_’, k_3_, k_4_, k_4,1_ k_5_, r_1_) and 4 outputs (VLP production, oligomer ratio, relative VLP production, VP40 budding ratio) are calculated.

The main output of interest was mature VLP production. Four key parameters were identified (k_4_, k_4,1_’, k_5_, r_1_) (Table S6) and they were all positively correlated to mature VLP production at all PS levels. Since r_1_ and k_5_ represented the ‘entry’ and ‘exit’ of VP40 in the system, their importance is expected. We noticed that k_2_ was significant when PS levels are low, suggesting VP40 membrane binding could be a rate-limiting step in low PS situations. Filament growth parameters (k_4_ and k_4,1_’) that drive the process of VLP production were also significantly correlated. However, it is surprising that k_4,1_’, a reverse rate constant had a positive correlation coefficient with VLP. Local sensitivity analysis revealed that increasing k_4,1_’ leaded to a significant increase in concentration of hexamer (Fig. S11). As a result, it created a strong hexamer pool that can overcome the increase in the reverse rate constant and facilitate filament growth. This counterintuitively suggested that the decrease of reverse constant rate for filament growth will disrupt VLP production.

PRCC results for relative VLP production among different PS groups identified k_1_, k_2_’, k_4,1_’ and k_5_ as significant in Ex3 model under low PS (14%) condition at 24h (Table S7). At 48h however, the significant parameters changed from k_1_ to k_4_ and r_1_ (Table S8). Here we focused on parameters having impact on longer term (48h) relative VLP production. Therefore, k_4_, k_2_’, k_4,1_’, k_5_ and r_1_ were positively correlated with both absolute VLP production number (Table S6) and relative VLP production at low PS level. This result suggested that targeting these parameters in combination with PS-targeted treatment may have synergistic effects in EBOV therapy.

Oligomer ratio is a metric of VP40 membrane binding and oligomerization. Since it represented the distribution of VP40 oligomers, any changes in the system might have an influence on it. As a result, almost all parameters were significantly correlated with the oligomer ratio (Table S9). PRCC results for VP40 budding ratio was similar to VLP production (Table S6, S10), as they both indicated the budding efficiency of VP40 VLPs.

Taken together, sensitivity analysis indicated that the VLP assembly process is robust to disruptions in stability of filaments by allowing levels of VP40 hexamers to compensate for differences in the reverse rate constant for filament growth (k_4,1_’). Results also indicated that parameter influences can be different for different PS levels, possibly identifying opportunities for combination therapy development.

## Discussion

The dynamics of VP40 oligomer assembly into EBOV matrix remained unknown. While membrane PS levels were known to affect VP40 assembly (35), the exact mechanisms of PS influence were still unclear. Using in vitro BSL-2 models (VP40 VLPs), integrated with computational modeling, we provided mechanistic insights into the VP40 assembly process. We developed the first intracellular model describing Ebola virus VP40 assembly and budding dynamics.

Our simulations indicated that the rate constant for initialization of VP40 oligomerization is different from that of continuous oligomerization, and that the growing filament structure stabilizes as it grows. While this stabilization effect has not been reported for VP40 filaments, our findings are consistent with the nucleation-elongation theory that has been established for other oligomers. It is known that for oligomerization of tubular or helical structures from single substrates, the initialization and elongation of the oligomerization are two different steps, and elongation is considered the faster step (55, 56) (e.g. microtubules (57, 58) and amyloid plaques (59–62)). Our results therefore suggested that this nucleation-elongation theory may also apply to VP40 oligomerization and could inform treatment strategies that target either nucleation or elongation steps.

Our prediction of preferential amplification of existing filaments (filament stabilization) is also consistent with structural studies showing VP40 to exist in a patchwork of assemblies at the plasma membrane inner leaflet (63, 64). These studies demonstrated that actin and VP40 diffused together and VP40 moved in a ballistic motion on these filaments in the absence of actin polymerization inhibitors (65). When actin polymerization was inhibited, VP40 exhibited constrained diffusion at the plasma membrane and VP40 enriched filaments emanating from the plasma membrane were significantly reduced (65). This transport mechanism of VP40 to sites of VLP assembly, in combination with our predicted nucleation-elongation kinetics of VP40 filament growth, would drive robust and effective VP40 assembly and budding.

One key finding of our simulations was that only when PS is directly affecting VLP budding can the experimentally observed longer term (48h) differences in relative VLP production be reproduced. We also needed to include the influence of PS on filament stabilization to reproduce the relative oligomer frequency difference among PS groups. These two mechanisms had not been previously reported. Our results indicated that PS might affect multiple steps in the viral budding process apart from the previously-identified VP40 membrane association (35). Thus, disruption of PS-VP40 interactions could be a promising drug target.

Sensitivity analysis indicated that production of VLP is also highly dependent on VP40 production, VP40 membrane association, filament assembly and VLP budding steps. Inhibiting those steps would decrease VLP production as well as relative VLP production at low PS level. Thus, while these steps could also be good targets alone (66), such as using graphene (67), a combination with PS targeting therapies such as fendiline (36, 68–70), staurosporine (71, 72) or bavituximab (73, 74) may have additional treatment efficacy.

Our model results showed fluctuations in oligomer concentration, oligomer ratio and mature VLP production for some parameter combinations. The fluctuations are caused by the dependence of the reverse rate constant 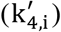 on filament size. Since this reverse rate constant decreases as the filament size increases, the transition of VP40 into smaller oligomers becomes out of balance with the ‘flow’ of VP40 into the larger oligomers. Since larger oligomers are more stable than smaller oligomers in filament growth, VP40 temporarily preferentially accumulates either in oligomers that are close to the size of a full filament or the hexamer pool. Further, the decrease in reverse rate constant 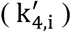 with filament size means that equilibrium between smaller oligomers is established faster than equilibrium between larger oligomers. This discontinuous size distribution of oligomer, plus the difference in equilibration time, together cause the wave-like patterns in filament growth.

Biological evidence also exists for fluctuations in elongating structures, for example in assembly of microtubules (75). Future experiments (e.g., real time imaging using total internal reflection microscopy as well as fluorescence correlation spectroscopy of VP40 at the plasma membrane interface and/or on supported lipid bilayers) can verify existence of fluctuating dynamics in VP40 assembly. The phenomenon suggests the possibility of noise in experimental data from single time points, especially at single cell levels.

As with all computational models, certain limitations apply to our model and analysis. We did not explicitly include transcription and translation processes, which may decrease our model-predicted time to steady state. The effects of diffusion or transportation are also not included in the model, and our rate constants thus represent effective rate constants. Our model currently does not include VP40 octamer rings, since it was not required for VLP production in previous studies (30, 32). Our model also successfully reproduces experimental data without VP40 octamers, which supports this prior conclusion. However, VP40 octamer is believed to play an unknown but important role in EBOV replication life cycle (30), which could be included in future models as more data emerge. Our model is based on and calibrated to the VP40 VLP system, and therefore any impacts on the other EBOV proteins will need to be progressively incorporated as we and others work to translate our findings to live EBOV dynamics.

Another limitation is that the experimental data are often collected from different conditions, time points and methods, leading to unavoidable differences between datasets. In some ways, this can be viewed as a strength as VP40 derived VLPs have been generated and used by many laboratories from different cell lines (HEK293, HUH7.5, Vero, A549, HeLa, CHO-K1 and others). Furthermore, the diversity in the data (Table S13) is exactly why computational models are a useful data integration tool. Overall, this study is focused on relative cost of model fits to determine the mechanisms needed to reproduce experimentally observed phenomena and trends.

In conclusion, we have built the first subcellular-level ODE-based model of the EBOV VP40 VLP system. We illustrated how the model can be qualitatively and quantitatively compared to data from *in vitro* VP40 VLP experimental studies. Our computational approach allowed complementary analysis that proposed that PS may have direct influence on VP40 filaments oligomerization, stabilization, and budding. Our combined experimental and computational approaches would enable further identification of key EBOV infection mechanisms and evaluation of treatment strategies and could be applied to similar viral systems.

## Experimental procedures

### ODE-based model construction

The structure of the hexamer-assembly-based model is summarized in Fig. 2. ODEs for the process are presented in equations (1)–(8). Total simulation time of the model is 48h.

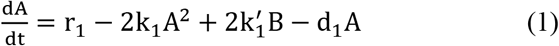

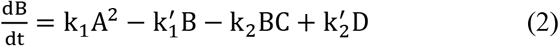

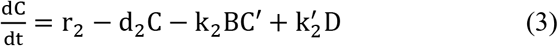

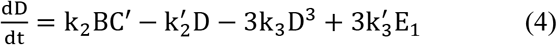

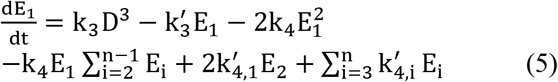

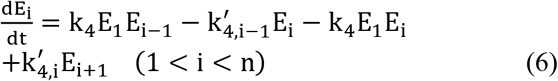

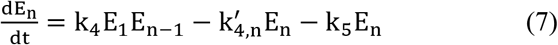

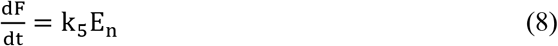

Initial conditions:

A(0) = 0
B(0) = 0
C(0) = 6.33 × 107 × PS(0)
D(0) = 0
E_i_(0) = 0 (1 ≤ i ≤ n)
F(0) = 0
PS(0) = 14%, 14.39%, 16.52%, 20%, 30% respectively

A: VP40 monomer in cytoplasm (nM).
B: VP40 dimer in cytoplasm (nM).
C: Total phosphatidylserine (nM).
C’: Phosphatidylserine available to interact with cytoplasmic VP40 dimer (nM, see equation (12)).
D: VP40 dimer on cell membrane (nM).
E(i): Developing matrix protein consists of i VP40 hexamers (nM).
i: Number of hexamers in developing filament.
n: Number of hexamers in a mature filament. n= 770 in our model.
F: Budded VLP (nM).
PS: Total phosphatidylserine (%).

PS level will be updated by the concentration of C through equation (9).

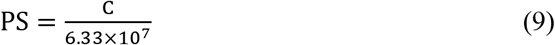

The dimer-assembly-based model excluded the assembly of hexamer as a separate process (Fig. S8), and ODEs for the process are shown in equation S(1)-S(7) (See supporting information).

### Influence of PS on VP40 dimer binding to membrane

Data from a surface plasmon resonance (SPR) experiment on VP40-PS affinity were curve fitted to derive an equation for K_D_ as well as available PS as a function of PS concentration in the membrane during the membrane binding process (36). K_D_ and PS available for the binding process were calculated through the following steps. We considered the following reversible binding reaction:

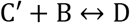

B: VP40 dimer in cytoplasm (nM).
C’: Phosphatidylserine available to interact with cytoplasmic VP40 dimer (nM).
D: VP40 dimer on cell membrane (nM). At steady state:

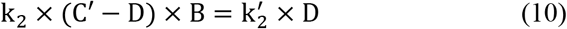

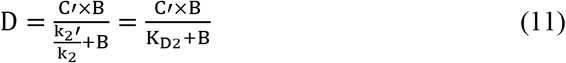
K_D2_: Equilibrium constant for VP40 dimer membrane binding (Table2).

K_D2_ and C’ were fitted through equation (11) using the SPR data (Table S11). Fitted values for K_D2_ and C’ at each PS level are included in Table S4.

Those SPR data were subsequently fitted into empirical equations (12) and (13) to enable us to calculate C and K_D2_ values at different PS levels in our simulation.

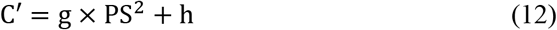

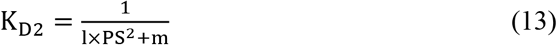

Fitted values of g, h, l and m are included in Table 2.

**Table 2.** Model parameters.

| Parameter | Value | Lower bound | Upper Bound |
| --- | --- | --- | --- |
| $k_1$ | Calibrated | $5e-4 /(\text{nM s})$ | $5e-2 /(\text{nM s})$ |
| $K_{D1}$ | 50 # | | |
| $k_1'$ | $k_1 \times K_{D1}$ | | |
| $k_2$ | $k_2' / K_{D2}$ | | |
| $K_{D2}$ | Calculated according to PS, l and m (Equation (13)). | | |
| $k_2'$ | Calibrated | $2.5e-6 /s$ | $2.5e-4 /s$ |
| $k_3$ | Calibrated <sup>*0</sup><br>Calibrated and calculated according to PS and x (Equation (21)) <sup>*1</sup> | $7e-7 /(\text{nM}^2 \text{ s})$ | $7e-5 /(\text{nM}^2 \text{ s})$ |
| $K_{D3}$ | 200 # | | |
| $k_3'$ | $k_3 \times K_{D3}$ | | |
| $k_4$ | Calibrated <sup>*0</sup><br>Calibrated and calculated according to PS and x (Equation (21)) <sup>*2</sup> | $1.5e-4 /(\text{nM s})$ | $1.5e-2 /(\text{nM s})$ |
| $k_{4,1}'$ | Calibrated | $7e-2 /s$ | $7 /s$ |
| $k_{4,i}'$ | Calculated according to PS, i, y, $t_{1-3}$ and $u_{1-3}$ (Equation (20)) | | |
| $k_5$ | Calibrated <sup>*0</sup><br>Calibrated and calculated according to PS and x (Equation (21)) <sup>*3</sup> | $7e-6 /s$ | $7e-4 /s$ |
| $r_1$ | Calibrated | $0.2 \text{ nM/s}$ | $20 \text{ nM/s}$ |
| $r_2$ | $d_2 \times C(0) (\text{nM})$. | | |
| $d_1$ | $2.25e-5$ (78) | | |
| $d_2$ | $2.5927e-5$ (79) | | |
| $n$ | 770 (80, 81) | | |
| $x$ | Calibrated <sup>*1,2,3</sup> | 0 | 3.333 |
| $y$ | Calibrated | 3 | 7 |
| $g$ | 35.16, fitted | | |
| $h$ | $1.479e+04$, fitted | | |
| $l$ | $9.196e-06$, fitted | | |
| $m$ | $4.523e-4$, fitted | | |
| $t_{1-3}$ | Fitted and calculated according to y | | |
| $u_{1-3}$ | Fitted and calculated according to y | | |

\*0: Baseline model. \*1, \*2, \*3: Ex1, Ex2 and Ex3 models. #: data from Dr. Stahelin (available on request).

### Influence of PS on filament stabilization

Filament stabilization was implemented by allowing the reverse rate constant to decrease as oligomerization increased until reaching a constant (non-zero) value. To account for PS influence on filament stabilization, the ratio of reverse rate constant of i^th^ oligomer to hexamer was defined as *f*(i, PS) in equation (14).

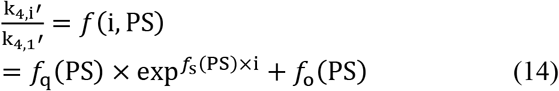

i: Number of hexamers in developing filament.

To include the influence of PS on this step, relative hexamer frequency data (transformed from the average of 3 replicates of oligomer frequency data) (36) was used to determine *f*_o_(PS), *f*_q_(PS) and *f*_s_(PS).

However, the frequency changing itself was not really the reverse rate constant for oligomerization. We only wished to have a trend of how k_4,i_’ decreased with oligomer size, not absolute values of how fast they decreased. Therefore, another parameter y was introduced to calculate a modified i in equation (15).

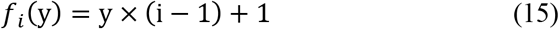

Combining equation (13) with (14) we obtain equation (15):

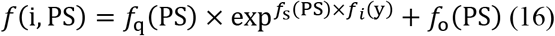

Values of *f*_o_(PS), *f*_q_(PS) and *f*_s_(PS) were then estimated by fitting equation (16) to experimental relative oligomer frequency data at the PS levels that were measured (Table S12). To estimate *f*_o_(PS), *f*_q_(PS) and *f*_s_(PS) at all PS levels, we defined their relationship with PS level by equation (16)–(18), where t_1~3_ and u_1~3_ were fitted using measured PS levels and corresponding values of *f*_o_(PS), *f*_q_(PS) and *f*_s_(PS) (Table 2).

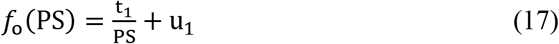

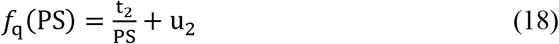

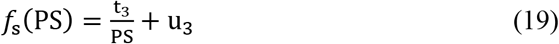

Combining equation (14) and (17–19), we get equation (20).

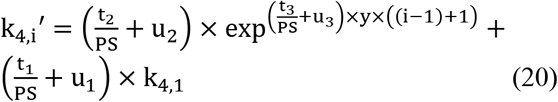

The value of parameter y would be calibrated during the procedure outlined in Table 2, and the estimation of t_1~3_ and u_1~3_ would be repeated for each calibration step. Finally we calculate k_4,i_’ by t_1~3_, u_1~3_, y, PS level and i through equation (20).

### Influence of PS on VP40 assembly and budding

Additional influences of PS on VP40 assembly and budding process were explored in three extended models using equation (21)

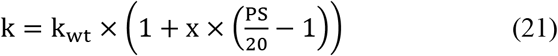

K_D2_: Equilibrium constant for VP40 dimer membrane binding (Table2).
k: involved parameter k_3_, k_4_ or k_5_ changing with PS level in Ex1, Ex2 or Ex3 separately.
k_wt_: involved parameter k_3_, k_4_ or k_5_ under 20% PS in Ex1, Ex2 or Ex3 separately.

Calibrated values can be found in Table S14-S17.

### Experimental data used for model calibration and validation

We have 5 types of data under 5 different PS level available for model calibration (Table 3).

- VP40 oligomer ratio from PSA-3 and PSA-3 with PS supplementation groups. The oligomer ratio is defined as the ratio of VP40 hexamers or larger oligomers relative to dimers and monomers (35).
- VLP production number from HEK293 cells at 24 hours (36) (Opinion from Dr. Stahelin).
- Relative oligomer frequency from HEK293, HEK293 treated with 1μM or 5μM of fendiline (36), which was transformed from N&B data and then averaged. Relative oligomer frequency is defined as the frequency of each VP40 oligomer, from membrane dimer to 42-mer, relative to the sum of oligomers (from membrane dimer to 42-mer). Concentration of larger oligomers was not experimentally detectable.
- Relative VLP production from HEK293, HEK293 treated with 1μM or 5μM of fendiline (36). Relative VLP production from PSA-3 group (35). Relative VLP production is defined as the amount of VLP in each group relative to wild type (WT) PS levels.
- VP40 budding ratio from HEK293 cell line (50). VP40 budding ratio is defined as the ratio of VP40 in VLPs relative to VP40 in cell. Details of data are included in Table S13.

**Table 3.** Experimental data.

|  | PS level | 14% | 14.39% | 16.52% | 20% | 30% |
| --- | --- | --- | --- | --- | --- | --- |
| | Experimental conditions | PSA-3 (genetic PS deficient) | HEK293 treated with 5μM fendiline | HEK293 treated with 1μM fendiline | HEK293 (WT) | PSA-3 supplied with additional PS |
| Data type | Oligomer ratio (24h) | √ |  |  |  | √ |
|  | VLP production (24h) |  |  |  | √ |  |
|  | Relative oligomer frequency (24h) |  | √ | √ | √ |  |
|  | Relative VLP production (24h) |  | √ | √ | √ |  |
|  | Relative VLP production (48h) | √ | √ | √ | √ |  |
|  | VP40 budding ratio (48h) |  |  |  | √ |  |

Validation data includes VP40 plasma membrane localization which is defined as the percentage of membrane VP40 from dimer to 42-mer in the detectable VP40s including cytoplasmic VP40s. The data is generated from live cell imaging experiments, performed at 8 hours and 24 hours post-transfection of WT EGFP-VP40 into HEK293 cells as previously described (36). HEK293 transfected cells were stained with Hoechst 3342 nuclear stain and a wheat germ agglutinin (WGA, Invitrogen Alexa Fluor 647) plasma membrane (PM) stain. Imaging was performed on a Nikon confocal microscope and image analysis was performed using Image J to determine the %PM localization for each time point. At least 24 cells were imaged per time point over three independent experiments on three different days.

### Parameter estimation and calibration

Model parameters are described in Table 2. The determination of parameters is outlined in Fig. 8. Since there were limited data available on the value of rate constants from direct measurements, we calibrated the rate constants through available experimental data from literature and our own work. The model was implemented in Matlab and solved using “ode15s”.

**Figure 8.**
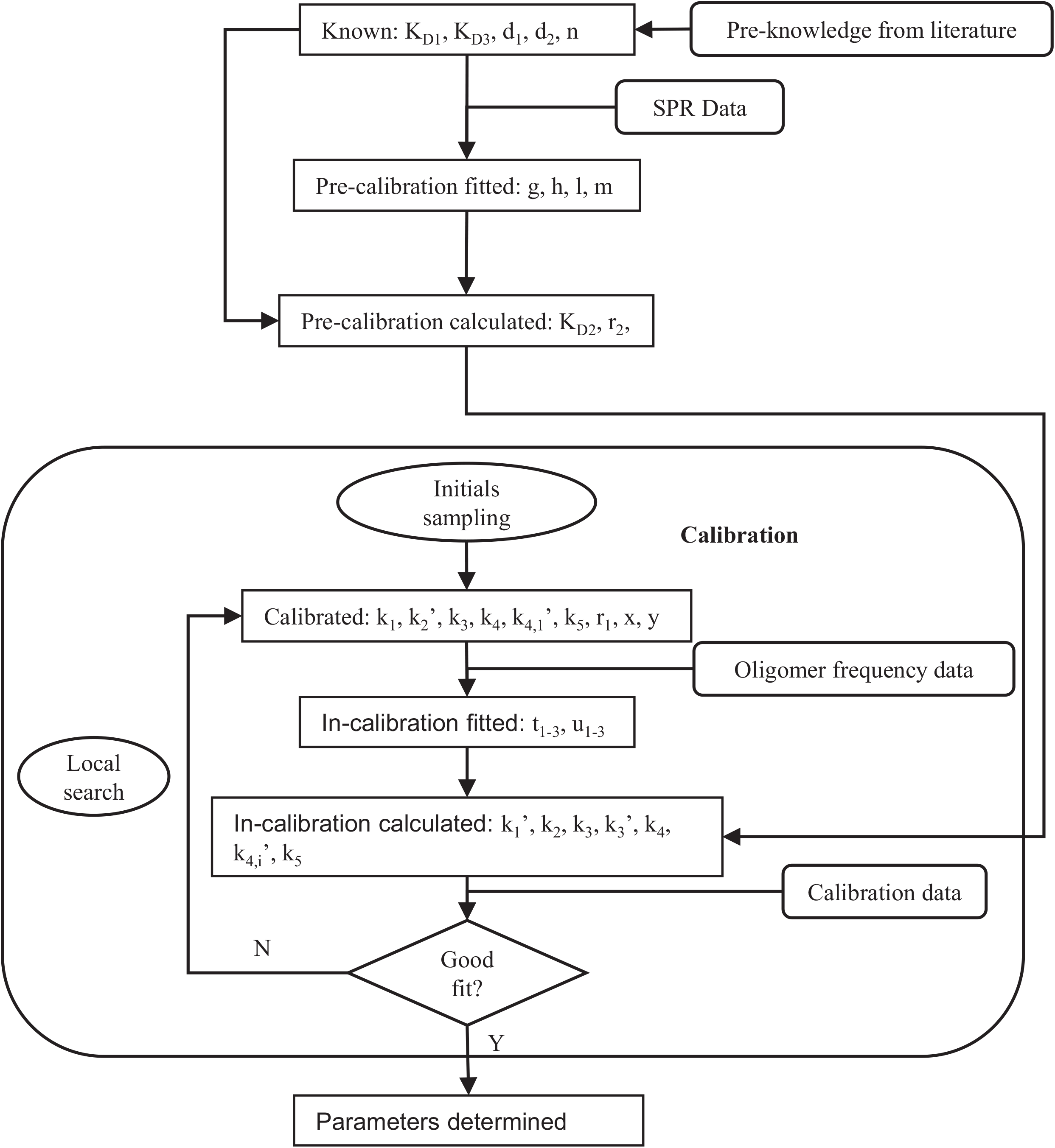
Parameterization process. All parameters can be organized into 6 categories and most of them are determined in the calibration cycle. The in-calibration calculations of k_3_, k_4_ and k_5_ are only conducted in Ex1, Ex2 and Ex3 models separately, and they will only undergo the first calibration process for the Baseline model.

We calibrated our model using the available data (Table 3) to identify parameters that minimize the cost calculated by equation (22).

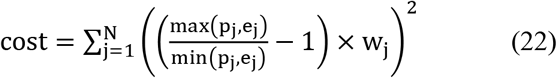

N: total number of experimental data points
e_j_: jth experiment data
p_j_: jth model prediction
w_j_: weight assigned to jth data point (Table S5)

Calibration was done using Matlab 2019b, where we used nonlinear least-squares solver (“lsqnonlin”) for optimization and Latin hypercube sampling (“lhsdesign”) for sampling initial guesses (76). The sampling for k_1_, k_2_’, k_3_, k_4_, k_4,1_’, k_5_ and r_1_ were in log-scale, while for x and y were linear scale. For each model, 50 initial guesses are generated. These initial guesses were used to initialize 50 independent optimizations and identify 50 parameter sets. Lower bounds and upper bounds used to sample initial guesses for each parameter are included in Table 2. The top 5 models (out of 50) with lowest cost function values were analyzed (Table S14-S17).

### Sensitivity analysis

Partial Rank Correlation Coefficient (PRCC) was used to perform global sensitivity analysis (77) to quantify the impact of k_1_, k_2_’, k_3_, k_4_, k_4,1_’, k_5_, r_1_, x and y on various model outputs, including: mature VLP, oligomer ratio, relative VLP production among PS groups and VP40 budding ratio. PRCC ranks each parameter and target output and calculates the partial correlation coefficient between them while taking other parameter variations into account. PRCC was done with Matlab 2019b, where we used partial correlation (“partialcorr”) for coefficient values calculation. Sampling range and method for k_1_, k_2_’, k_3_, k_4_, k_4,1_’ and k_5_ was the same as calibration (Table 2). Empirical parameters ‘x’ and ‘y’ were fixed at 3.2 and 5 respectively in order to focus our analysis on physiological parameters. Each parameter was sampled 500 times, and PRCCs were calculated for each model separately (baseline model and three extended models). P-values lower than 0.05 were regarded as significant.

Local sensitivity analysis was performed by fixing all parameters at values from the top fit of each model and varying the parameter of interest within 2 orders of magnitude.

## Data Availability

The majority of the data are contained within the article and supporting information. For data not included within the article, data can be shared by contacting the corresponding author.

The code can be obtained from DOI: 10.5281/zenodo.5106604.

## Supporting information

This article contains supporting information.

## Acknowledgements

We would like to acknowledge the Purdue Pharmacy Live Cell Imaging Facility for confocal imaging (NIH OD027034 to R.V.S.)

## Funding and additional information

These studies were supported by the NIH/NIAID (AI081077) to R.V.S., the NIH/NIGMS to M.L.H. (T32GM075762), the Indiana CTSI (UL1TR001108) to R.V.S and E.P and the Frederick N. Andrews Fellowship to X.L.

The content is solely the responsibility of the authors and does not necessarily represent the official views of the National Institutes of Health.

## Abbreviations and nomenclature

PS: phosphatidylserine
VLP: virus-like particle
EBOV: Ebola virus
EVD: Ebola virus disease
MEURI: Monitored Emergency Use of Unregistered and Investigational Interventions
ODE: ordinary differential equation
SPR: surface plasmon resonance
VP40: viral protein 40.

