## Supporting information for "Mechanisms of phosphatidylserine influence on viral production: a computational model of Ebola virus matrix protein assembly"

**ODE-based dimer assembly model construction:**

Equation (S1)-(S7) represent the VP40 assembly pathway in the dimer model which assumes VP40 dimer to be the building block of filaments.

$$\frac{dA}{dt} = r_1 - 2k_1A^2 + 2k_1'B - d_1A \qquad (S1)$$

$$\frac{dB}{dt} = k_1A^2 - k_1'B - k_2BC' + k_2'D_1 \qquad (S2)$$

$$\frac{dC}{dt} = r_2 - d_2C - k_2BC' + k_2'D_1 \qquad (S3)$$

$$\frac{dD_1}{dt} = k_2BC - k_2'D_1 - 2k_4{D_1}^2 - k_4D_1\sum_{i=2}^{n-1} D_i + 2k_{4,1}'D_2 + \sum_{i=3}^{n} k_{4,i}'\,D_i \qquad (S4)$$

$$\frac{dD_i}{dt} = k_4D_1D_{i-1} - k_{4,i-1}'D_i - k_4D_1D_i + k_{4,i}'D_{i+1} \quad (1 < i < n) \qquad (S5)$$

$$\frac{dD_n}{dt} = k_4D_1D_{n-1} - k_{4,n}'D_n - k_5D_n \qquad (S6)$$

$$\frac{dF}{dt} = k_5D_n \qquad (S7)$$

initial conditions:

$A(0) = 0$

$B(0) = 0$

$C(0) = 6.33 \times 10^7 \times PS(0)$

$D_i(0) = 0 \quad (1 \leq i \leq n)$

$F(0) = 0$

$PS(0) = 14\%, 14.39\%, 16.52\%, 20\%, 30\%$ respectively

A: VP40 monomer in cytoplasm (nM).

B: VP40 dimer in cytoplasm (nM).

C: Total phosphatidylserine (nM).

C‘: Phosphatidylserine available to interact with cytoplasmic VP40 dimer (nM).

$D_i$: Developing matrix protein consists of i VP40 dimers (nM).

i: Number of dimers in developing filament.

n: Number of dimers in a mature filament. n= 2310 in our model (nM).

F: Budded VLP (nM).

PS: Total phosphatidylserine (%).

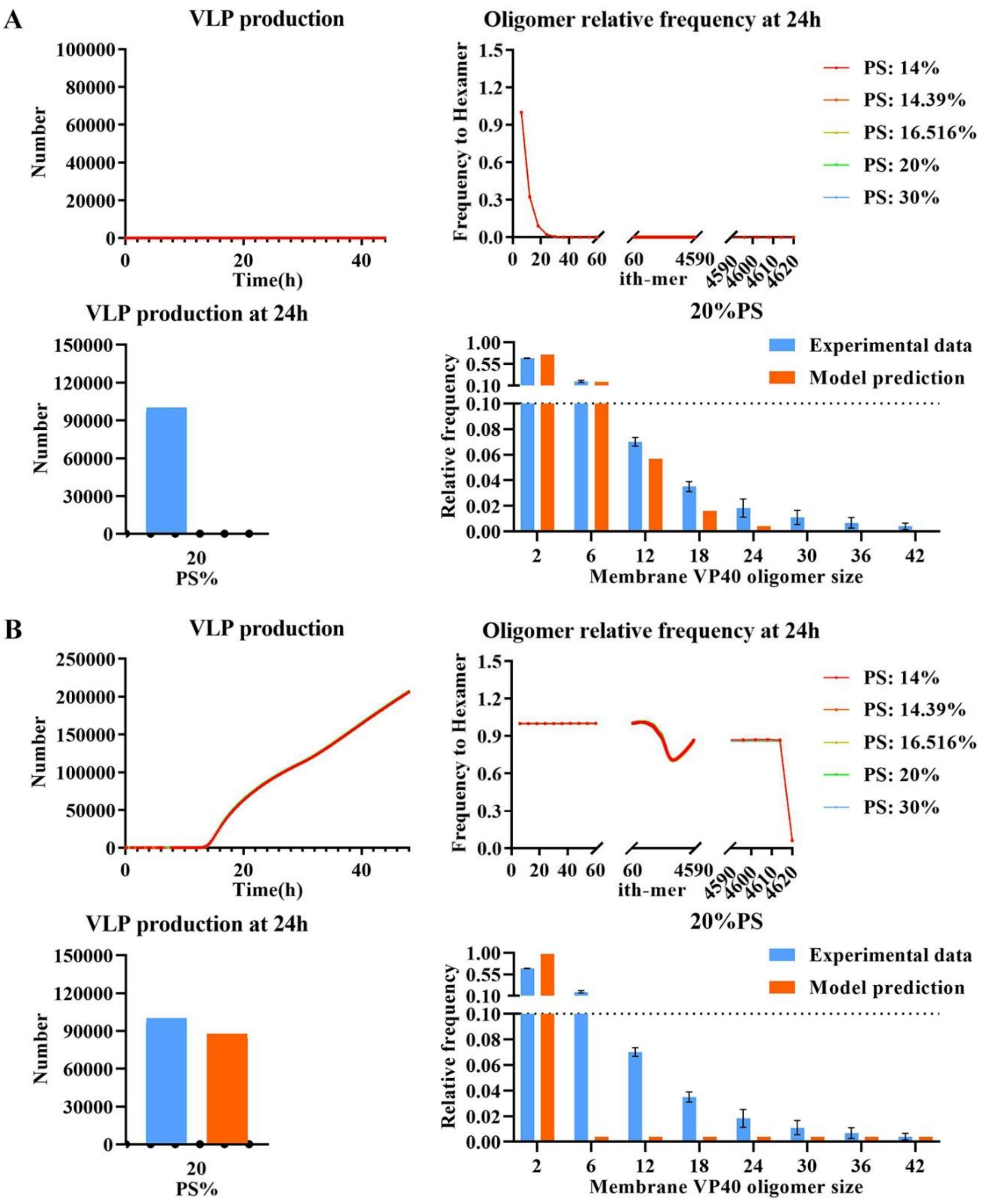


**Figure 1. The influence of PS on the EBOV budding process.** PS is known to influence VP40 dimer membrane association, oligomer profile and VLP production. However, whether the observed phenomena can be explained by the VP40 dimer association alone remains unclear. Error bar indicates the SEM. Sample sizes of each experimental data are shown in Table S13.

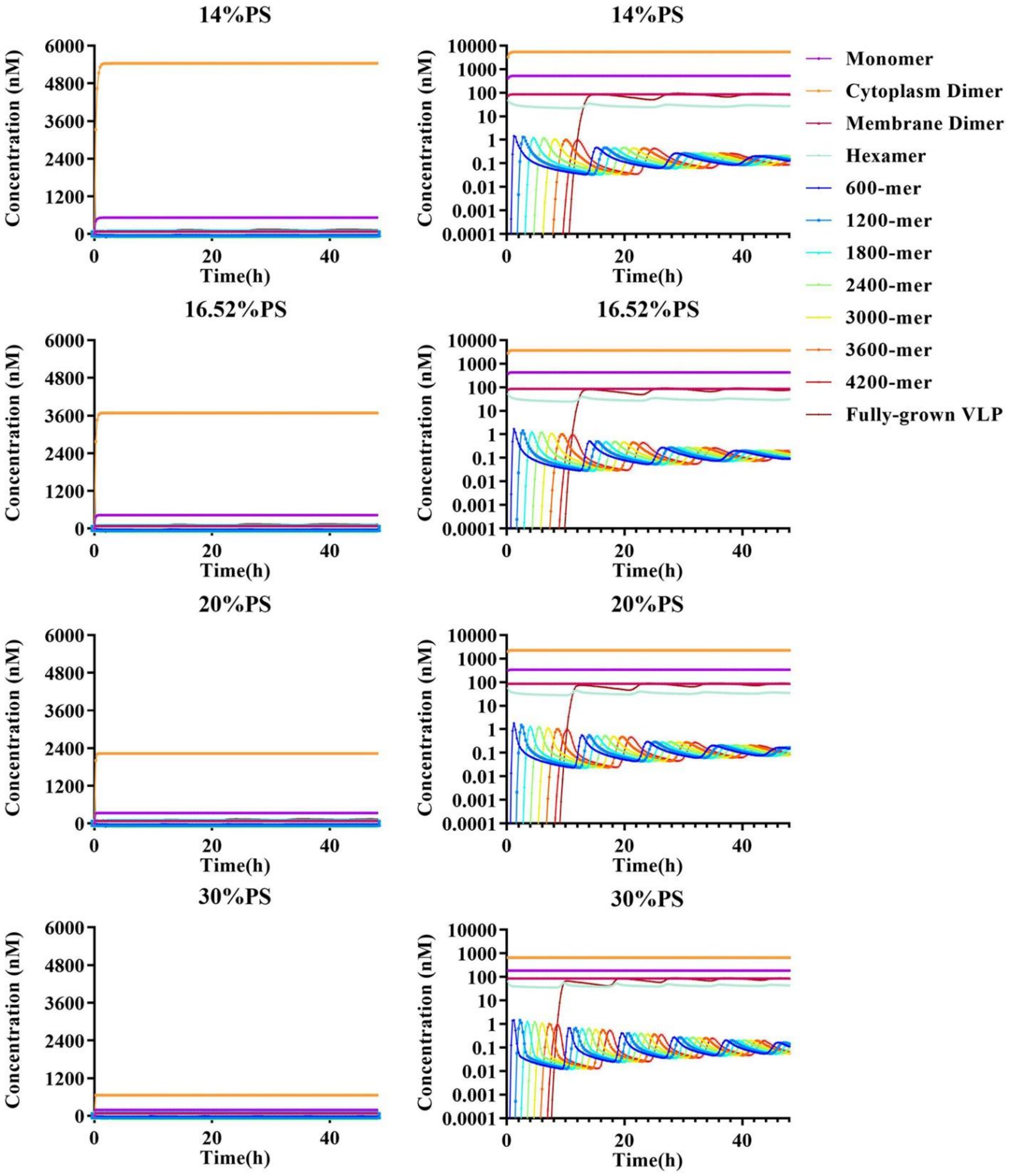


**Figure S2. Time course of VP40 monomer and oligomers of best fit in basic model.** PS level ranges from 14% to 30%. Left panels are on linear-scale and right panels are on log-scale.

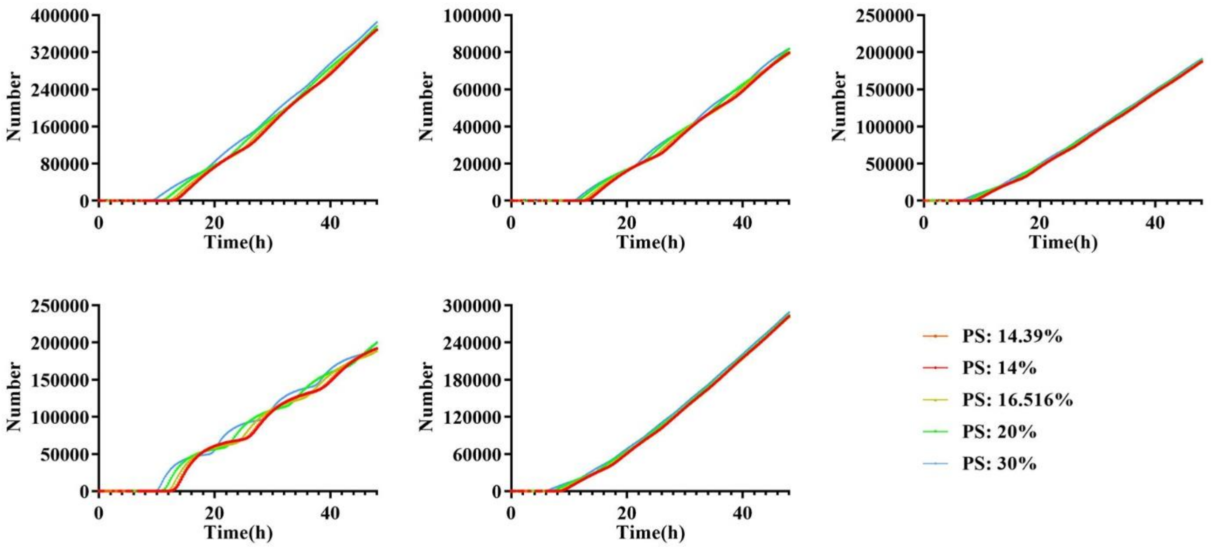


**Figure S3. VLP production dynamic of baseline model from top5 fittings.** VLP production for each PS concentration under baseline model are overlapping.

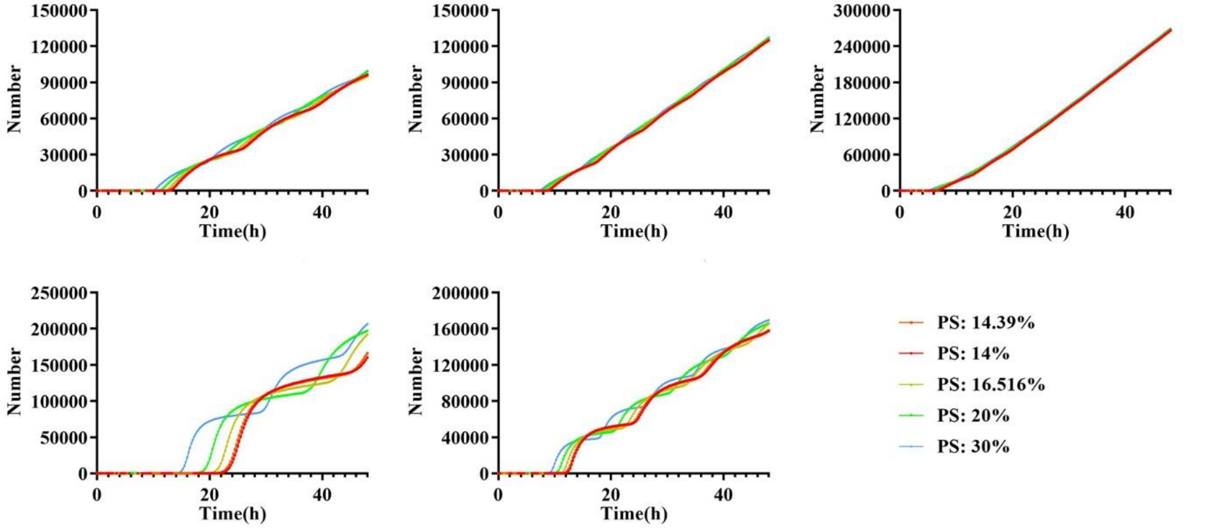


**Figure S4. VLP production dynamic of Ex1 model from Top5 fittings.** VLP production for each PS concentration under Ex1 model are overlapping.

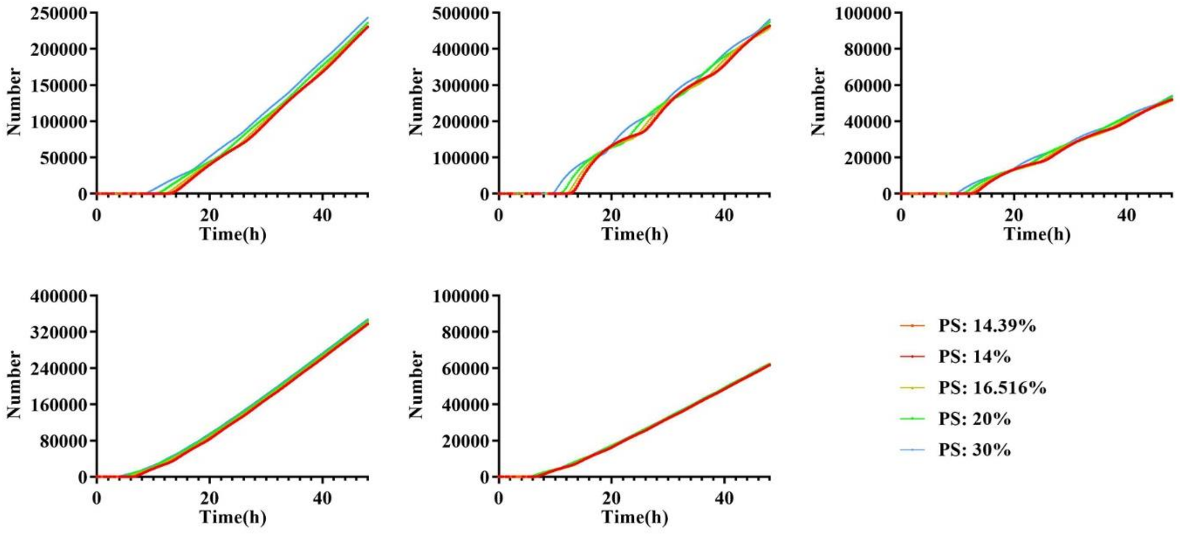


**Figure S5. VLP production dynamic of Ex2 model from Top5 fittings.** VLP production for each PS concentration under Ex2 model are overlapping.

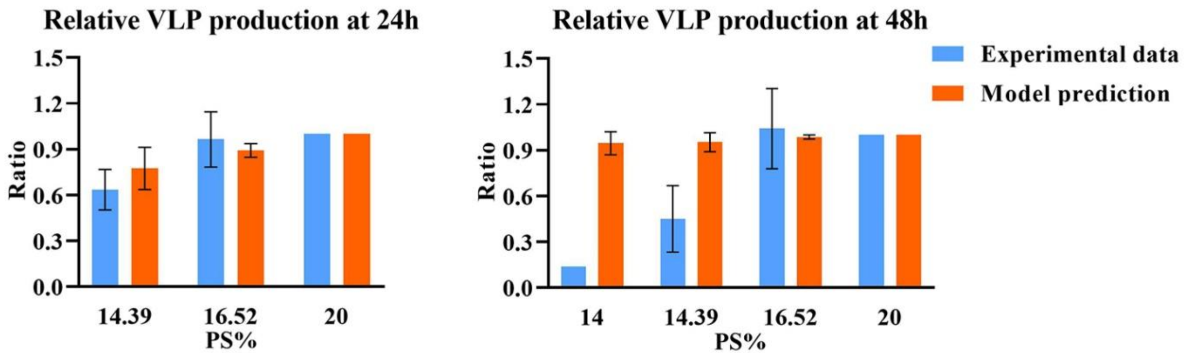


**Figure S6. Relative VLP production at 24 (left) and 48h (right) of Ex1 model from top5 fittings.** No obvious difference can be observed in different PS groups, especially for 48h. Error bar indicates the SEM. Simulation data represents top 5 fits. Sample sizes of each experimental data are shown in Table S13.

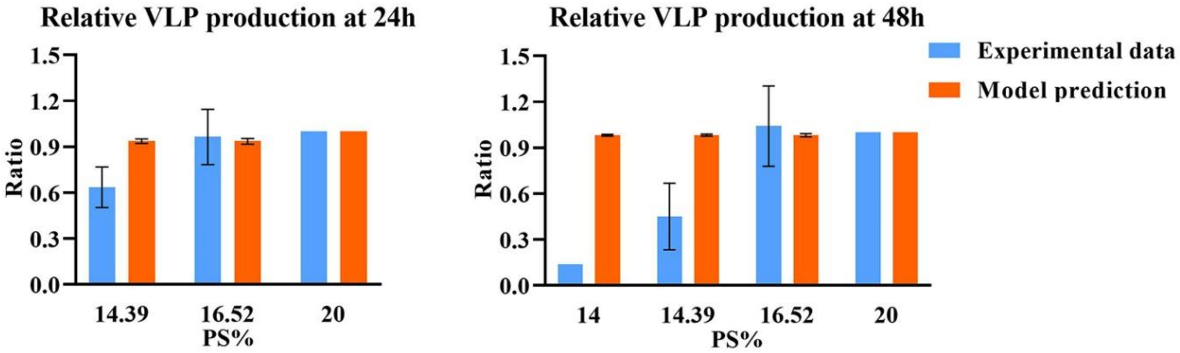


**Figure S7. Relative VLP production 24 (left) and 48h (right) of extended model 2 from top5 fittings.** No obvious difference can be observed in different PS groups, especially for 48h. Error bar indicates the SEM. Simulation data represents top 5 fits. Sample sizes of each experimental data are shown in Table S13.

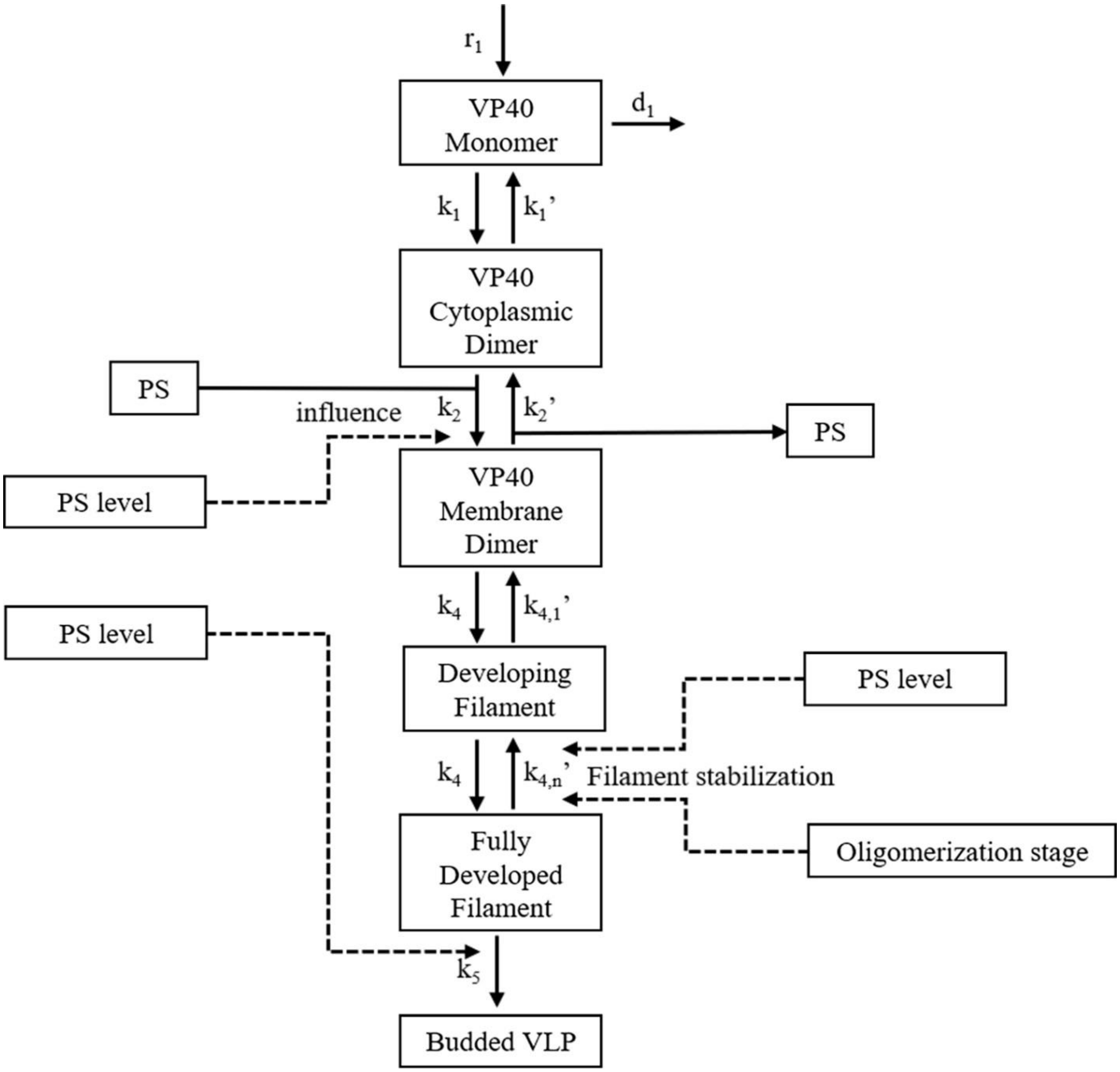


**Figure S8. Scheme of VP40 dimer assembly model.** The dimer assembly model is similar to hexamer assembly model. VP40 hexamer is deleted in this model, and filaments are directly built from VP40 membrane dimer. The model includes the influence of PS on VLP budding rate.

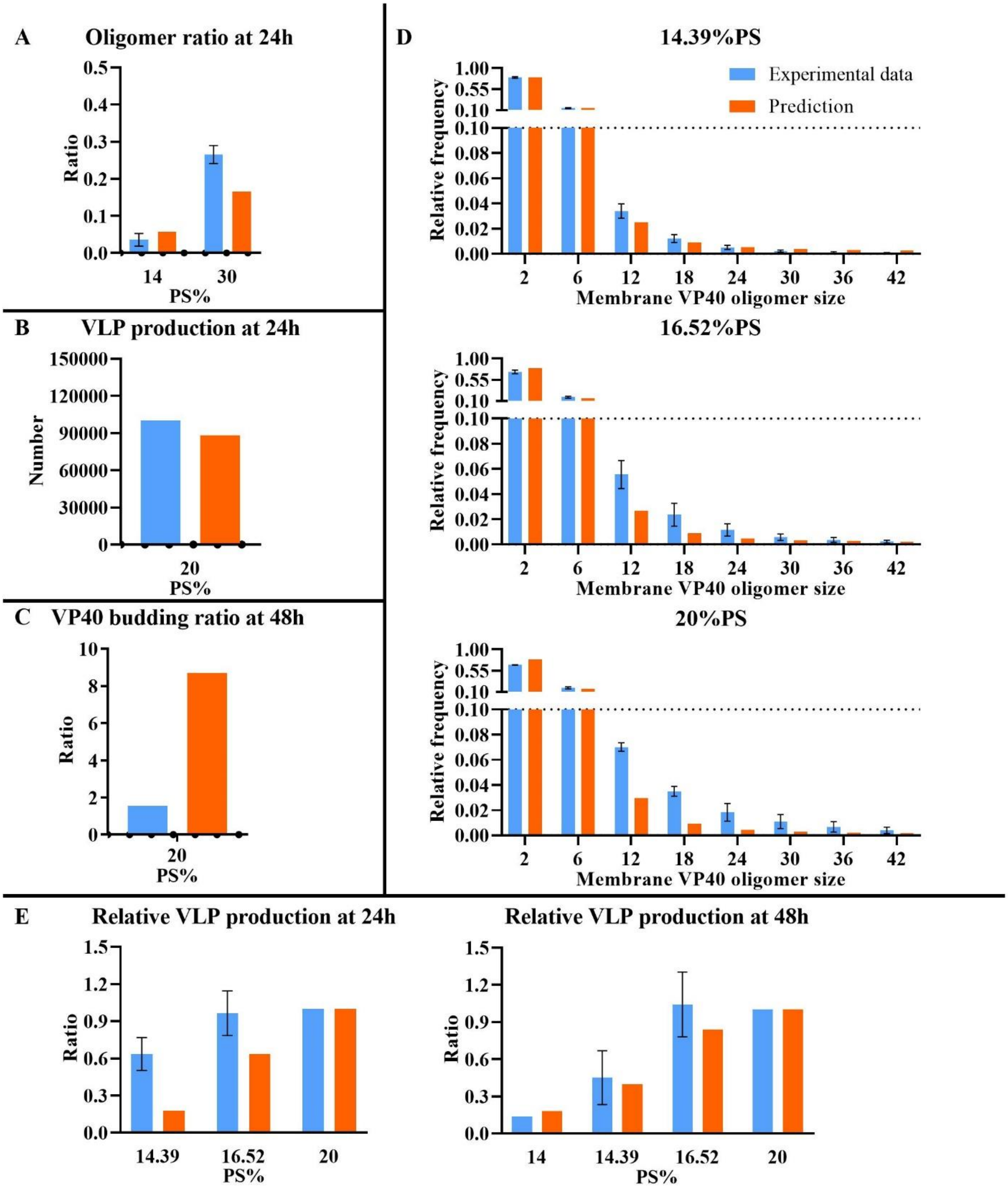


**Figure S9. Comparison between prediction and experiment data in dimer-based model (Ex3).** (A) Oligomer ratio. (B) VLP production. (C) VP40 budding ratio. (D) Oligomer frequency. (E) Relative VLP production. Error bar indicates the SEM. Sample sizes of each experimental data are shown in Table S13.

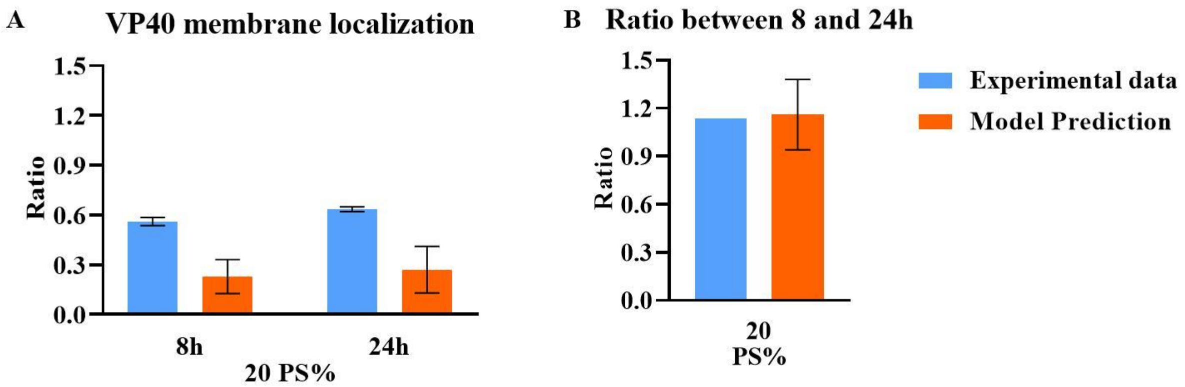


**Figure S10. Verification of Ex3 through VP40 membrane localization.** (A) Prediction of Ex3 model and experimental data on VP40 membrane localization at both 8h and 24 h. (B) A ratio of VP40 membrane localization at 24h and 8h. Error bar indicates the SEM. Simulation data represents top 5 fits. Sample sizes of each experimental data are shown in Table S13.

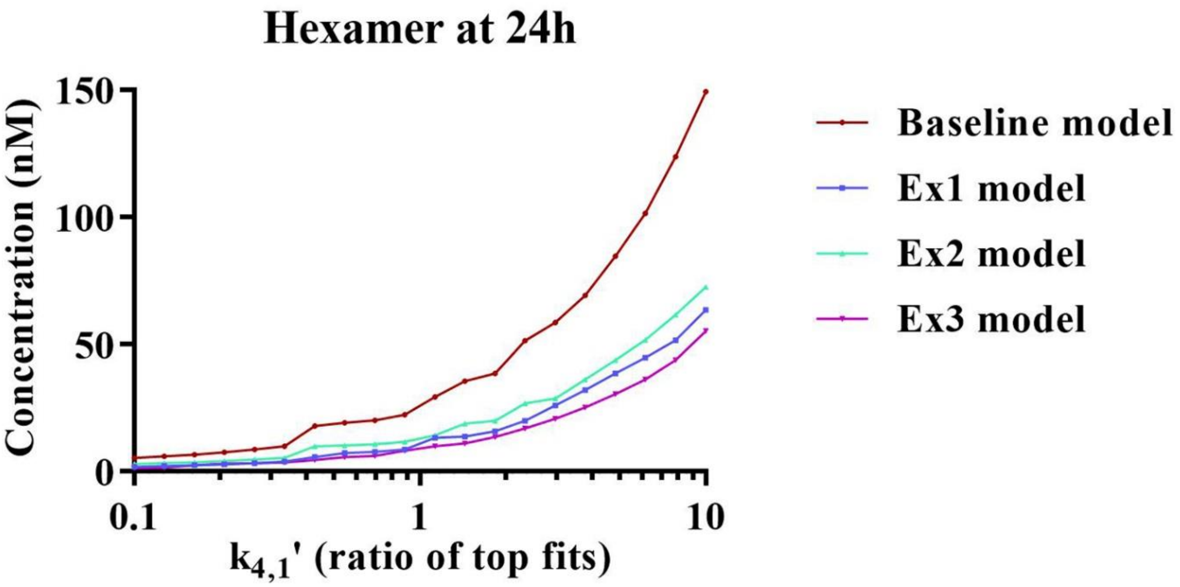


**Figure S11. Local sensitivity analysis of hexamer to $k_{4,1}$.** In each of the model, hexamer concentration is positively correlated to $k_{4,1}$.

**Table S1. Top 5 lowest calibration cost for all models.**

| Top | Baseline | Ex1 | Ex2 | Ex3 |
|---|---|---|---|---|
| 1 | 40.189 | 44.0296382 | 43.82747726 | 15.65277585 |
| 2 | 41.89715 | 45.03716865 | 46.95113682 | 20.02522637 |
| 3 | 44.81008 | 45.92155152 | 46.9863814 | 22.39429639 |
| 4 | 46.44327 | 48.67798537 | 47.20208397 | 23.04378811 |
| 5 | 49.58851 | 51.82399249 | 47.68961903 | 28.7278176 |
| **Average** | **44.5856** | **47.09806724** | **46.53133969** | **21.96878086** |

**Table S2. One-way ANOVA for cost among models.**

| ANOVA | | | | | | |
|---|---|---|---|---|---|---|
| Source of Variation | SS | df | MS | F | **P-value** | F crit |
| Between Groups | 2195.926 | 3 | 731.9752 | 60.01585 | **6.33E-09** | 3.238872 |
| Within Groups | 195.1418 | 16 | 12.19636 | | | |
| Total | 2391.067 | 19 | | | | |

**Table S3. LSD for cost between models.**

| | t | p-value | significant |
|---|---|---|---|
| baseline VS Ex1 | -1.1375 | 0.864 | N |
| baseline VS Ex2 | -0.8809 | 0.804 | N |
| baseline VS Ex3 | 10.2397 | 9.87E-09 | Y |
| Ex1 VS Ex2 | -0.2566 | 0.600 | N |
| Ex1 VS Ex3 | 11.1206 | 3.07E-09 | Y |
| Ex2 VS Ex3 | 11.3772 | 2.22E-09 | Y |

**Table S4. Fitted values of C and $K_{D2}$**

| **PS (%)** | **C (nM)** | **$K_{D2}$ (nM)** |
|---|---|---|
| 1 | $1.761*10^4$ | 2158 |
| 11 | $1.535*10^4$ | 639.9 |
| 22 | $3.273*10^4$ | 203.9 |

**Table S5. Weight of calibration.**

| Data | w |
|---|---|
| VLP production number | 0.5[1] |
| VP40 oligomer ratio | 0.8[2] |
| Relative oligomer frequency | 1 |
| Relative VLP production | 1 |
| VP40 budding ratio | 1 |

1: Weight lowered because it is an estimated value.
2: Weight lowered because the interpretation of this data may not be accurate.

**Table S6. PRCC for VLP production.**
See “Supporting_information_PRCC.xlsx”

**Table S7. PRCC for relative VLP production at 24h.**
See “Supporting_information_PRCC.xlsx”

**Table S8. PRCC for relative VLP production at 48h.**
See “Supporting_information_PRCC.xlsx”

**Table S9. PRCC for Oligomer ratio.**
See “Supporting_information_PRCC.xlsx”

**Table S10. PRCC for VP40 budding ratio.**
See “Supporting_information_PRCC.xlsx”

**Table S11. SPR data.**
See “Supporting_information_data.xlsx”

**Table S12. Transformed relative oligomer frequency data for filament stabilization.**
See “Supporting_information_data.xlsx”

**Table S13. Data for calibration.**
See “Supporting_information_data.xlsx”

**Table S14. Calibration result for baseline model.**
See “Supporting_information_calibration.xlsx”

**Table S15. Calibration result for Ex1 model.**
See “Supporting_information_calibration.xlsx”

**Table S16. Calibration result for Ex2 model.**
See “Supporting_information_calibration.xlsx”

**Table S17. Calibration result for Ex3 model.**
See “Supporting_information_calibration.xlsx”

**Table S18. VP40 membrane localization at 8 h.**
See “Supporting_information_VP40_Membrane_localization.xlsx”

**Table S19. VP40 membrane localization at 24 h.**
See “Supporting_information_VP40_Membrane_localization.xlsx”
